# Upregulation of distinct miRNAs in SARS-CoV-2 infected individuals: A differential signature of circulating miRNAs

**DOI:** 10.64898/2026.08.02.742360

**Authors:** Íris Terezinha Santos de Santana Silva, Sandra Rocha Gadelha, Hllytchaikra Ferraz Fehlberg, Fabrício Barbosa Ferreira, Mylene de Melo Silva, Rachel Passos Rezende, George Rego Albuquerque, Ana Paula Melo Mariano, João Carlos Teixeira Dias, Galileu Barbosa Costa, Carla Martins Kaneto

## Abstract

COVID-19 exhibits a broad clinical spectrum, from asymptomatic to severe cases, underscoring the need for molecular biomarkers to support surveillance and early intervention. Here, we profiled circulating microRNAs (miRNAs) in plasma samples from individuals with asymptomatic/mild COVID-19 and uninfected controls. Seven miRNAs were significantly upregulated in infected patients (miR-126, miR-146b-5p, miR-223-5p, miR-144-3p, miR-22, miR-146a, and miR-30c). ROC curve analysis revealed heterogeneous diagnostic performance: miR-30c achieved the highest overall discriminatory accuracy (AUC = 0.771) with maximum sensitivity (100.0%), while miR-126 provided the highest specificity (100.0%, AUC = 0.763). Other miRNAs, including miR-146b-5p, miR-223-5p, miR-146a, miR-144-3p, and miR-22, showed intermediate accuracy (AUCs 0.684–0.719), whereas miR-21-5p and miR-155 displayed limited discriminatory power (AUCs 0.606 and 0.517, respectively). Predictive interaction network analysis indicated that the upregulated miRNAs target key immune-related genes (CXCL12, IRAK1, TRAF6, STAT1, JAK1, NOTCH1, SMAD4, and BCL2L11), and functional enrichment revealed convergence with transcriptomic profiles from SARS-CoV-2-infected Calu-3 cells, including FOXO3, JAK2, STAT1, and SIRT1. Collectively, these findings point out for potential miRNA signatures associated with mild, non-hospitalized COVID-19 in a predominantly vaccinated cohort but requiring further investigation as molecular markers of early host responses in larger, independent, and clinically diverse cohorts.

**Importance:** COVID-19 produces a wide range of outcomes, from no symptoms to severe illness, and clinicians still lack simple molecular tools to help identify infections or anticipate a patient’s course early in the disease. miRNAs are small molecules circulating in blood that help control gene activity, and because their levels change during infection, they can be considered potential biomarkers candidates for blood-based tests. Here, we measured nine circulating miRNAs in individuals with mild COVID-19 and in uninfected individuals, thus finding that seven of them were consistently higher in infected patients, with some distinguishing the two groups quite well. This study also links these miRNAs to genes involved in the immune response against SARS-CoV-2. Our findings, generated in a modestly sized cohort, support further investigation of blood-based microRNA panels as potential candidates to help identify infection and clarify early host responses to SARS-CoV-2, pending validation in larger and more clinically diverse cohorts.

## INTRODUCTION

Coronavirus disease 2019 (COVID-19), caused by severe acute respiratory syndrome coronavirus 2 (SARS-CoV-2), has triggered a global public health crisis since its emergence in late 2019 (1). SARS-CoV-2 is a highly transmissible RNA virus belonging to the *Coronaviridae* family, characterized by its ability to infect human epithelial cells through binding to the angiotensin-converting enzyme 2 (ACE2) receptor (2, 3). SARS-CoV-2 is primarily transmitted via respiratory droplets and direct contact, with an incubation period ranging from 2 to 14 days (4).

The clinical manifestations of COVID-19 have varied widely since the beginning of the pandemic, ranging from asymptomatic or mild flu-like symptoms to severe pneumonia, acute respiratory distress syndrome (ARDS), and multiorgan complication (4). While most individuals experience mild disease, a subset develops severe outcomes due to an exaggerated inflammatory response, characterized by cytokine storm and immune dysregulation. Advanced age, comorbidities such as cardiovascular disease, diabetes, and obesity, as well as genetic and immunological factors, have been associated with worse clinical outcomes (1, 5). Given this clinical variability, understanding the host–pathogen interactions underlying COVID-19 severity remains essential to improving early diagnosis, prognosis, and therapeutic strategies.

The immune response to SARS-CoV-2 plays a pivotal role in disease progression, balancing viral clearance with tissue damage. Innate immunity is activated upon viral entry, leading to the production of interferons (IFNs) and pro-inflammatory cytokines that shape the adaptive immune response. However, in severe cases, dysregulated immune activation results in hyperinflammation, endothelial dysfunction, and thrombotic complications (6, 7). Significant advances have been made in the diagnosis and prevention of COVID-19. RT-qPCR has become the gold standard for SARS-CoV-2 detection, providing high sensitivity and specificity (7, 8), while rapid antigen tests played a critical role in large-scale screening and outbreak control, and serological assays were employed to assess past infections and population-level immunity (9–11). Also importantly, the rapid development and deployment of vaccines drastically altered the pandemic trajectory. Several mRNA-based, viral vector, and inactivated virus vaccines were approved and widely distributed, demonstrating high efficacy in reducing severe disease and mortality (8, 11).

Despite these advances in COVID-19 diagnostics, identifying molecular biomarkers capable of predicting disease progression and distinguishing different outcomes remains a critical research priority (12). In this context, microRNAs (miRNAs) have emerged as promising biomolecules due to their role as essential post-transcriptional regulators of gene expression, playing a crucial role in immune modulation, inflammation, and antiviral defense (12, 13). These small non-coding RNAs regulate the expression of multiple target genes by promoting mRNA degradation or inhibiting translation, thereby influencing various biological processes, including host responses to viral infections (13). In the setting of COVID-19, alterations in miRNA expression profiles have been reported in both mild and severe cases, suggesting their potential involvement in disease pathogenesis and immune dysregulation (14–16). Moreover, miRNA signatures may provide insights into long-term disease complications, such as post-acute sequelae of COVID-19 or long COVID (17, 18).

Despite the growing body of evidence supporting the relevance of miRNAs as promising biomarkers for early detection and prognosis of SARS-CoV-2 infection, further studies are needed to validate their utility in clinical settings. That way, identifying miRNA expression patterns in individuals with different outcomes may help to refine predictive models for disease progression (16). Furthermore, analysis of potential target genes may provide insights into understanding pathogenesis and future paths to disease modulations. In this study, we aimed to analyze the differential expression of miRNAs in patients with mild COVID-19, as an initial step toward identifying potential candidates as molecular biomarkers of SARS-CoV-2 infection that could be further investigated in larger, independent cohorts spanning a broader range of disease severity.

## RESULTS

In the case group, 13 were female (52.0%) and 12 were male (48.0%), whereas in the control group, 20 were female (80.0%) and 5 were male (20.0%), with no statistically significant difference in gender and age distribution between groups (p = 0.072 and p = 0.924, respectively) (Table 1). Besides, all patients in the case group were non-hospitalized and presented with at least one mild clinical symptom, such as: cough, sore throat, chest pain, rhinorrhea, and anosmia or ageusia (loss of smell or taste).

**TABLE 1.**
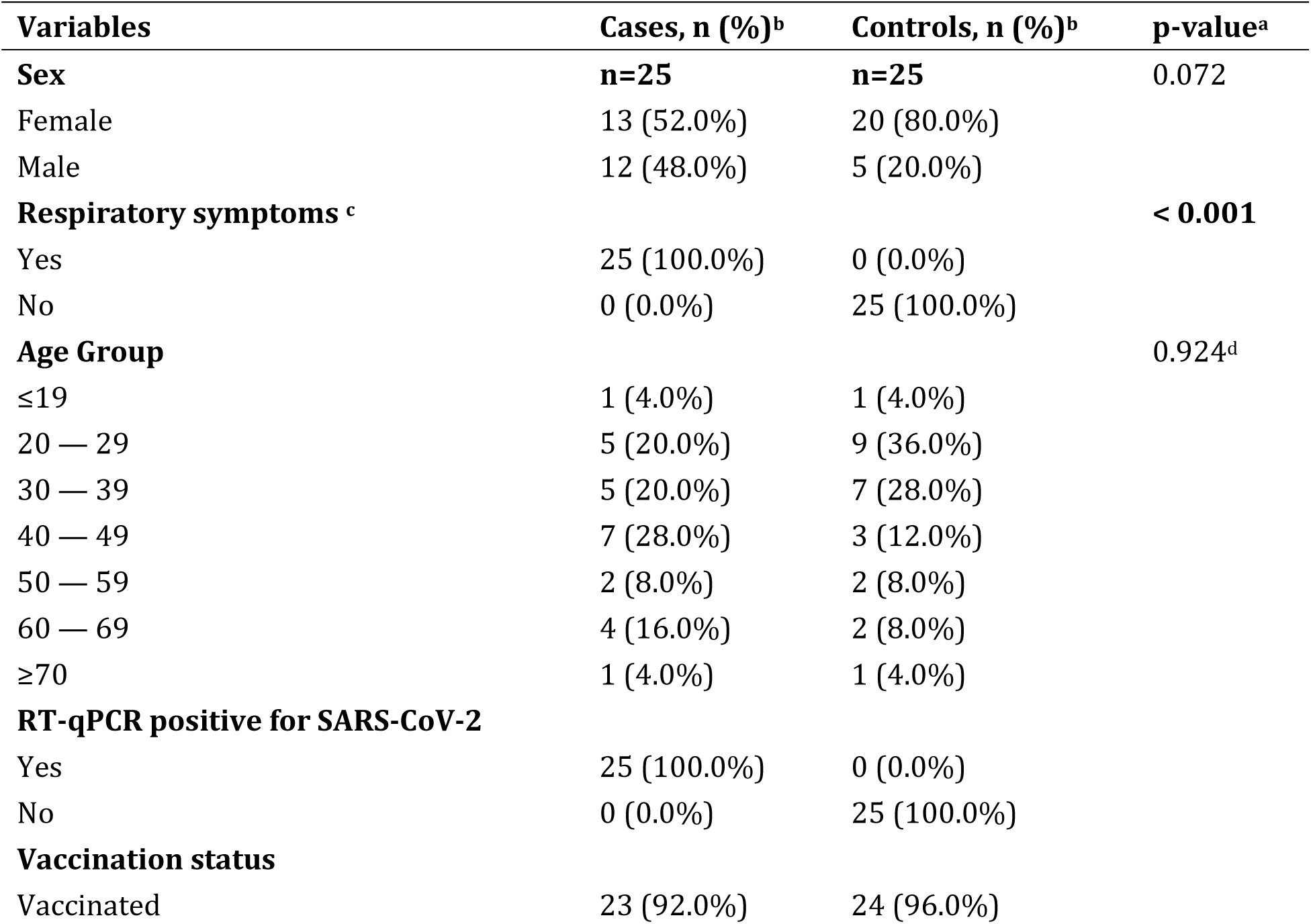
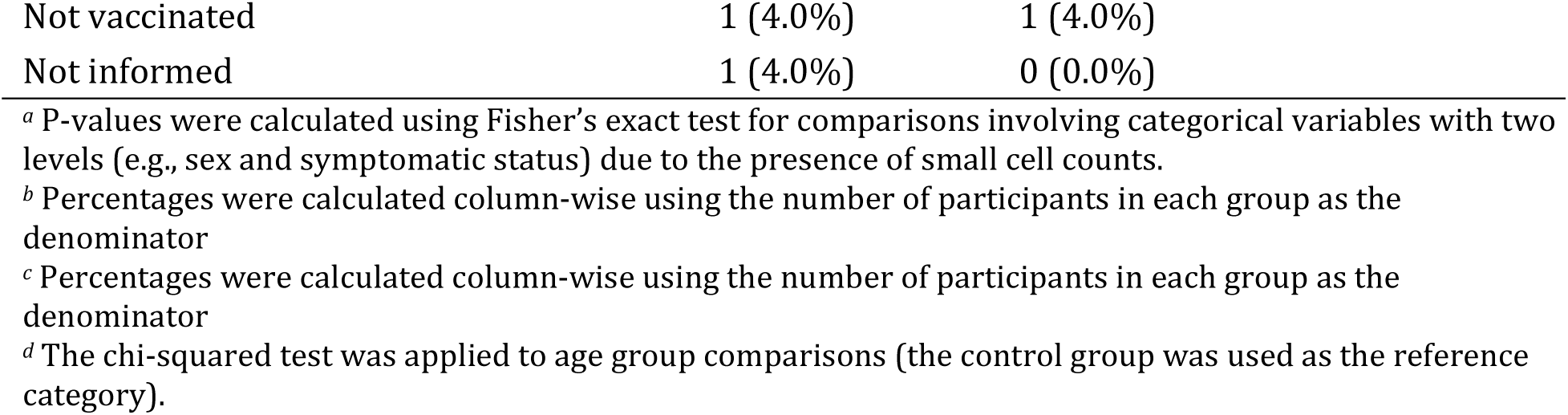
Clinical and demographic characteristics of enrolled participants by group (cases vs controls).

| Variables | Cases, n (%)<sup>b</sup> | Controls, n (%)<sup>b</sup> | p-value<sup>a</sup> |
| --- | --- | --- | --- |
| **Sex** | **n=25** | **n=25** | 0.072 |
| Female | 13 (52.0%) | 20 (80.0%) |  |
| Male | 12 (48.0%) | 5 (20.0%) |  |
| **Respiratory symptoms <sup>c</sup>** |  |  | **< 0.001** |
| Yes | 25 (100.0%) | 0 (0.0%) |  |
| No | 0 (0.0%) | 25 (100.0%) |  |
| **Age Group** |  |  | 0.924<sup>d</sup> |
| ≤19 | 1 (4.0%) | 1 (4.0%) |  |
| 20 — 29 | 5 (20.0%) | 9 (36.0%) |  |
| 30 — 39 | 5 (20.0%) | 7 (28.0%) |  |
| 40 — 49 | 7 (28.0%) | 3 (12.0%) |  |
| 50 — 59 | 2 (8.0%) | 2 (8.0%) |  |
| 60 — 69 | 4 (16.0%) | 2 (8.0%) |  |
| ≥70 | 1 (4.0%) | 1 (4.0%) |  |
| **RT-qPCR positive for SARS-CoV-2** |  |  |  |
| Yes | 25 (100.0%) | 0 (0.0%) |  |
| No | 0 (0.0%) | 25 (100.0%) |  |
| **Vaccination status** |  |  |  |
| Vaccinated | 23 (92.0%) | 24 (96.0%) |  |

|  |  |  |
| --- | --- | --- |
| Not vaccinated | 1 (4.0%) | 1 (4.0%) |
| Not informed | 1 (4.0%) | 0 (0.0%) |

<sup>a</sup> P-values were calculated using Fisher's exact test for comparisons involving categorical variables with two levels (e.g., sex and symptomatic status) due to the presence of small cell counts.
<sup>b</sup> Percentages were calculated column-wise using the number of participants in each group as the denominator
<sup>c</sup> Percentages were calculated column-wise using the number of participants in each group as the denominator
<sup>d</sup> The chi-squared test was applied to age group comparisons (the control group was used as the reference category).

The analysis of relative miRNA expressions in samples from cases vs controls revealed a significant upregulation of multiple miRNAs (Fig. 1). Specifically, a statistically significant upregulation was observed for miR-146b-5p (p < 0.05; Fig. 1A), miR-126 (p < 0.01; Fig. 1B), miR-223-5p (p < 0.05; Fig. 1C), miR-146a (p < 0.05; Fig. 1D), miR-30c (p < 0.01; Fig. 1E), miR-144-3p (p < 0.05; Fig. 1F), and miR-22 (p < 0.05; Fig. 1G) in the patients with mild COVID-19. The relative expression levels (expressed as 2^−ΔCt) for each of these miRNAs were notably higher in the case group, indicating robust positive regulation. Expression variability was observed in both groups, but the differences in medians (and interquartile ranges, when visible) were consistent with the statistical significance.

**FIG 1.**
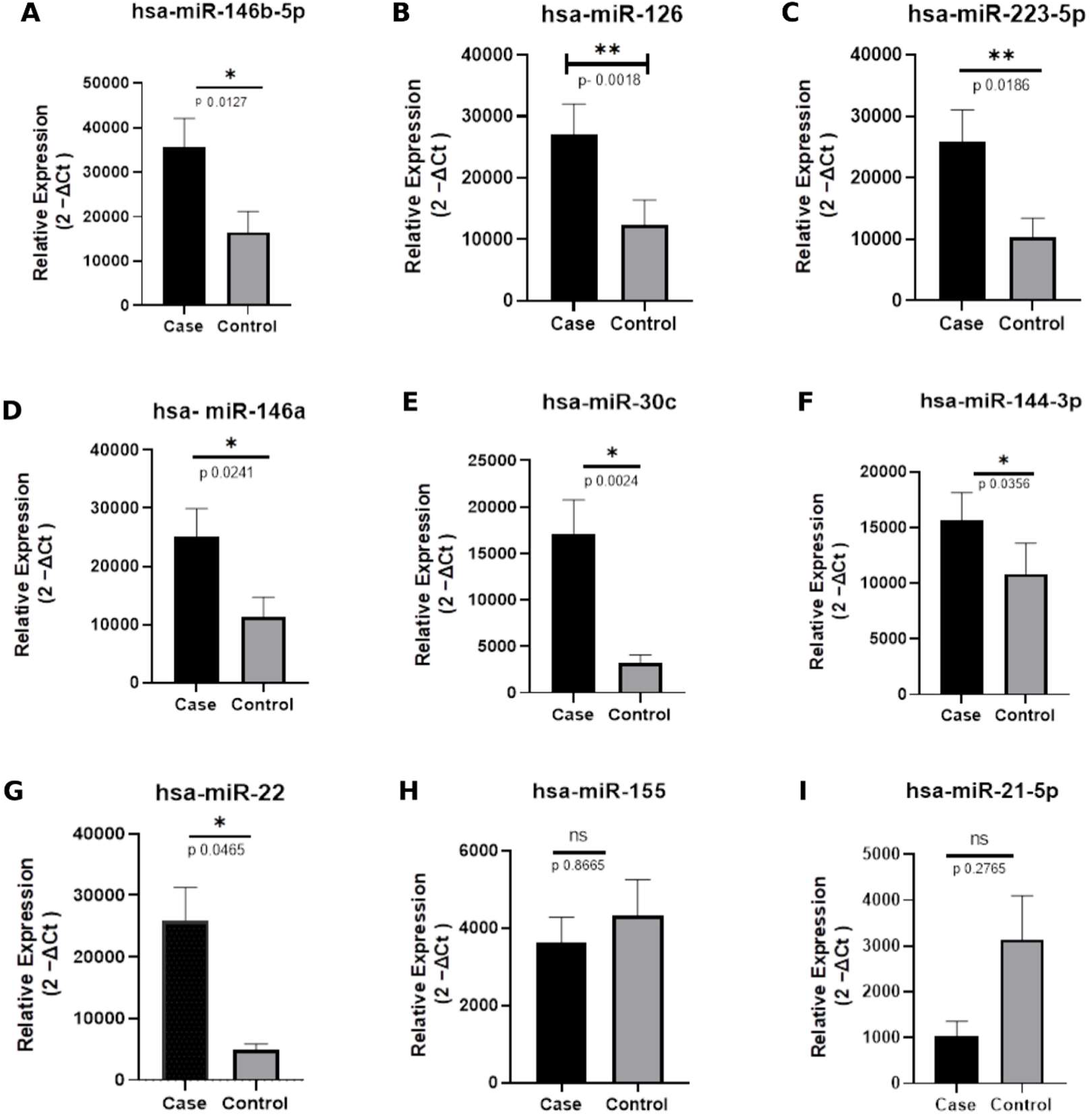
Relative expression of miRNAs in patients with mild COVID-19 (Case) and control subjects (Control). Scatter plots represent the relative expression (2−ΔCt) of specific miRNAs. Horizontal bars indicate the median and quartiles. Asterisks denote statistical significance: p < 0.05, p < 0.01.

To evaluate the potential use of these miRNAs as biomarkers for symptomatic/mild COVID-19, ROC curve analyses were performed (Fig. 2). The ROC curves demonstrated that most of the upregulated miRNAs exhibited good to excellent ability to distinguish between patients with mild COVID-19 and healthy individuals. Among them, miR-146b-5p achieved balanced diagnostic parameters with an AUC of 0.719 (95% CI: 0.566–0.871; p = 0.0133; Fig. 2A). At its Youden-optimal cut-off (< 17428), it showed a sensitivity of 75.0% and a specificity of 66.7%. miR-126 also displayed strong performance with an AUC of 0.763 (95% CI: 0.615–0.912; p = 0.0022; Fig. 2B). At its Youden-optimal cut-off (< 1489), it achieved high specificity (100.0%) and moderate sensitivity (59.1%). In turn, miR-223-5p exhibited an AUC of 0.704 (95% CI: 0.553–0.856; p = 0.0191; Fig. 2C). At its Youden-optimal cut-off (< 35067), it displayed high sensitivity (95.2%) but limited specificity (41.7%).

**FIG 2.**
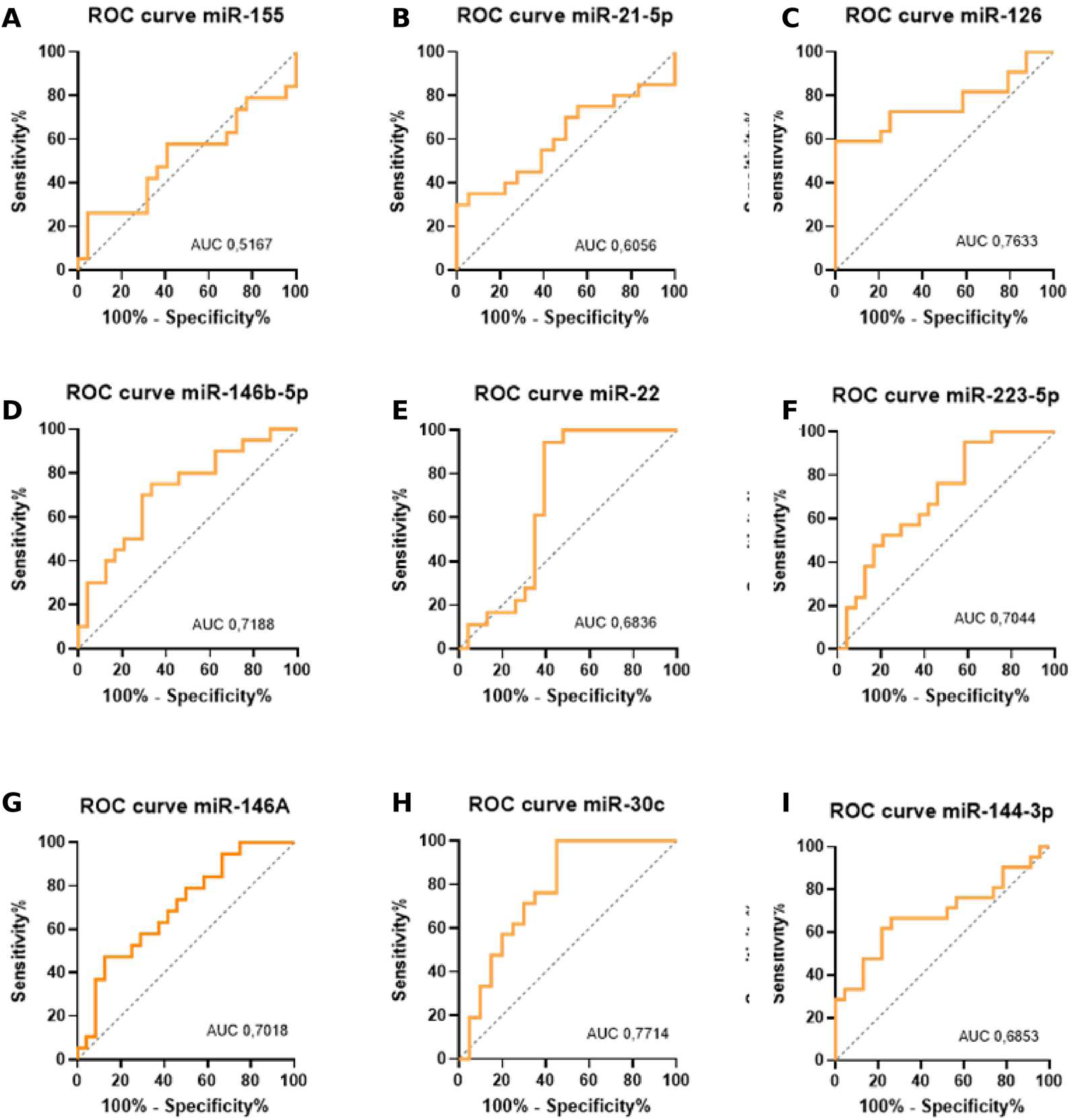
miRNAs selected as biomarkers of mild COVID-19. Each plot represents the sensitivity (%) vs. 100-specificity (%) for a specific miRNA, with the dotted line indicating the random chance (AUC = 0,5). Area Under the Curve (AUC) values are presented for each miRNA.

Regarding miR-146a, it demonstrated an AUC of 0.702 (95% CI: 0.546–0.858; p = 0.0245; Fig. 2D). At its Youden-optimal cut-off (< 1733), it presented a sensitivity of 47.4% and a specificity of 87.5%. miR-30c showed the highest overall performance, with an AUC of 0.771 (95% CI: 0.622–0.921; p = 0.0029; Fig. 2E). At its Youden-optimal cut-off (< 14486), it exhibited a sensitivity of 100.0% and a specificity of 55.0%. miR-144-3p and miR-22-3p also showed moderate performance, with AUCs of 0.685 (95% CI: 0.522–0.849; p = 0.0355; Fig. 2F) and 0.684 (95% CI: 0.511–0.856; p = 0.0459; Fig. 2G), respectively. At their Youden-optimal cut-offs, miR-144-3p exhibited a sensitivity of 66.7% and a specificity of 73.9% (< 8141), while miR-22-3p showed a sensitivity of 94.4% and a specificity of 60.9% (< 10015).

In contrast, miR-155 and miR-21-5p presented lower diagnostic performance, with AUCs of 0.517 (95% CI: 0.331–0.703; p = 0.8548; Fig. 2H) and 0.606 (95% CI: 0.422–0.789; p = 0.2666; Fig. 2I), respectively. At their Youden-optimal cut-offs, miR-155 showed a sensitivity of 26.3% and a specificity of 95.5% (> 7930), whereas miR-21-5p displayed a sensitivity of 30.0% and a specificity of 100.0% (> 5545). Despite these high specificities, their low AUCs and higher p-values indicate limited overall discriminatory capacity.

Additionally, an interaction network analysis based on target prediction data (retrieved from miRWalk) revealed potential interactions between several of the upregulated miRNAs and genes involved in immunological and cellular processes relevant to COVID-19 (Fig. 3). Specifically, hsa-miR-146a was predicted to interact with IRAK1 and TRAF6, both key mediators of toll-like receptor and NF-κB signaling. hsa-miR-146b-5p was also linked to STAT1, IRAK1, and TRAF6, suggesting convergent regulation of inflammatory pathways. hsa-miR-126-3p showed predicted interactions with CXCL12, DNMT1, and SMAD4. hsa-miR-30c-5p was associated with NOTCH1, BCL2L11, CXCL12, and DNMT1, implicating roles in apoptosis and immune modulation. hsa-miR-144-3p was predicted to regulate NOTCH1, CXCL12, and SMAD4, while hsa-miR-223-5p was linked to IL6R and BCL2L11. Furthermore, hsa-miR-22-3p was predicted to interact with SIRT1, FOXO3, PTEN, and BCL2L11, which are key regulators of cellular stress resistance, antioxidant responses, immune homeostasis, and apoptosis, respectively.

**FIG 3.**
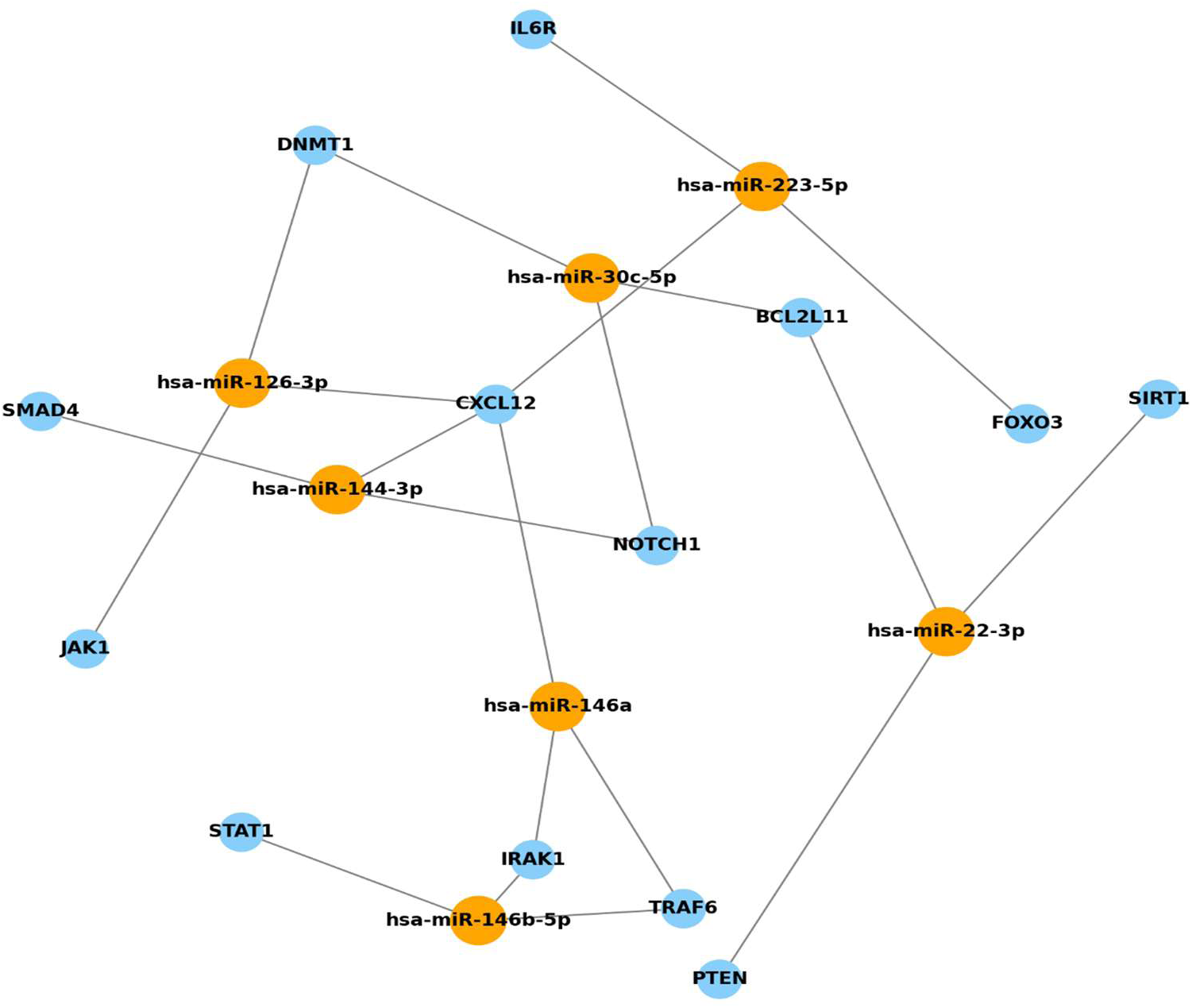
Network of interactions emphasizing miRNAs and shared genes. Orange nodes represent the identified up-regulated miRNAs. Blue nodes represent the genes predicted to target these miRNAs, based on interaction prediction data (mirWalk). The lines connect the miRNAs to their respective target genes.

Predictive functional enrichment analysis performed with the Metascape platform identified significant associations between the target genes of the differentially expressed miRNAs and several pathological conditions (Fig. 3A). The top categories included idiopathic pulmonary arterial hypertension, neuralgia, subarachnoid hemorrhage, and middle cerebral artery occlusion, as well as hematological malignancies such as precursor T-cell lymphoblastic leukemia-lymphoma, diffuse large B-cell lymphoma, low-grade B-cell malignancy, and cutaneous T-cell lymphoma. Additional enriched terms included vascular diseases, head and neck carcinoma, tubulointerstitial fibrosis, immune suppression, and myocardial reperfusion injury.

Finally, transcriptomic overlap analysis (Fig. 4B) revealed a significant convergence between the miRNA-regulated genes and gene sets previously described as upregulated in Calu-3 lung epithelial cells at 12 hours and 24 hours post-SARS-CoV-2 infection (19). This overlap included genes involved in type I interferon signaling, cytokine-mediated responses, and apoptosis regulation. Four predicted targets identified in this study, FOXO3, JAK2, STAT1, and SIRT1, were also found among the genes upregulated in SARS-CoV-2-infected Calu-3 cells at 12 hours (COVID038) and 24 hours (COVID040), according to the curated COVID-19 gene sets in Metascape (Figure 4B).

**FIG 4.**
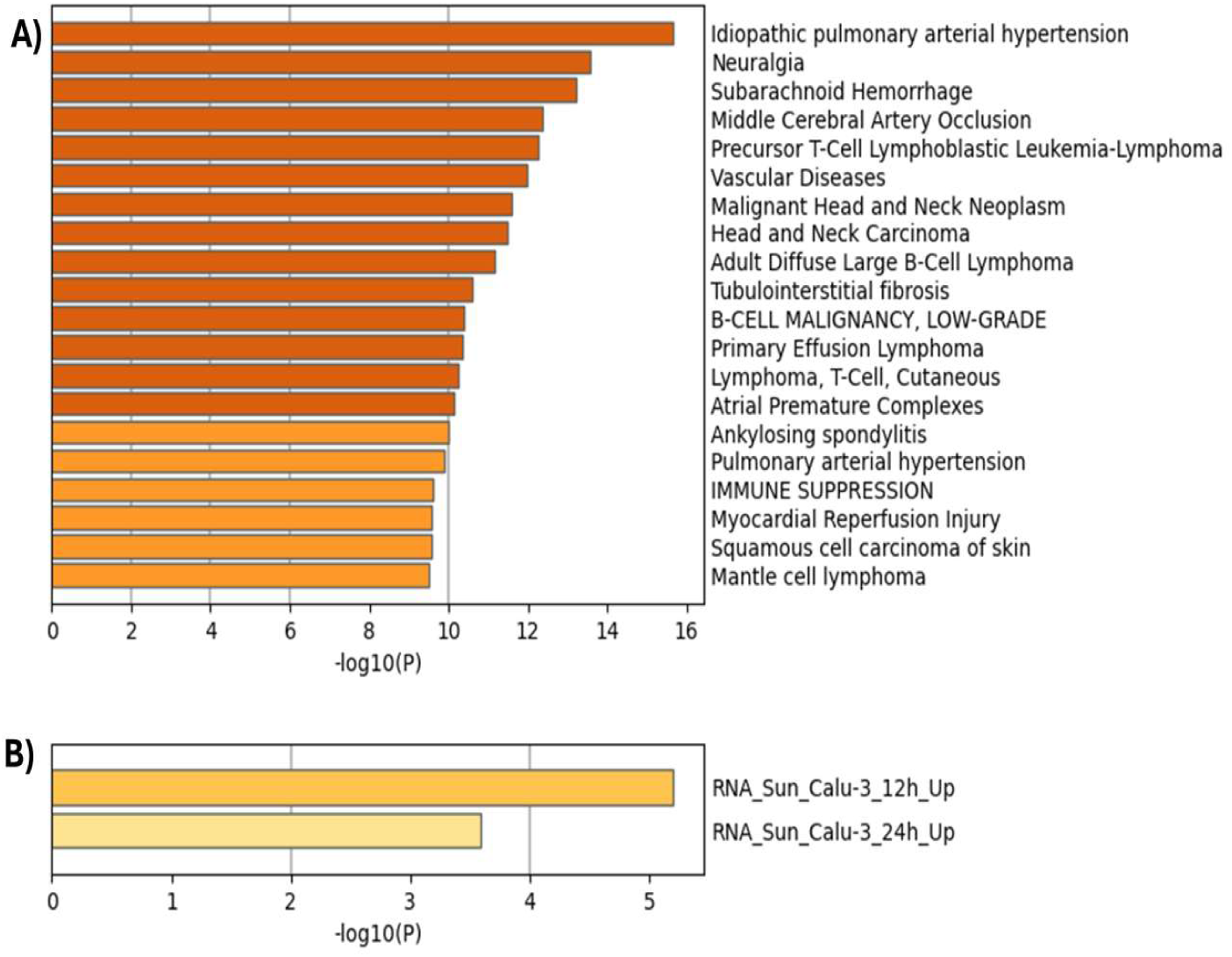
Enrichment analysis of disease-associated gene sets and transcriptomic signatures related to SARS-CoV-2 infection. (**A**) Top 20 diseases significantly associated with target genes predicted for the differentially expressed miRNAs (DisGeNET database). (**B**) Gene expression profiles upregulated in Calu-3 cells infected with SARS-CoV-2 at 12 h and 24 h post-infection (RNA-Seq dataset: Sun et al.). Bars represent –log₁₀(P) values from enrichment analysis. Data was generated using Metascape.

## DISCUSSION

This study investigated the expression profile of miRNAs in patients with mild COVID-19, identifying an upregulated miRNA signature potentially involved in the pathogenesis and in the host immune response. Notably, none of the detected miRNAs exhibited significantly downregulated expression, suggesting that the transcriptional response in mild COVID-19 is predominantly characterized by upregulation of specific miRNAs with regulatory functions, such as inflammation control, tissue repair, and modulation of apoptosis.

As for the upregulated miRNA, miR-146a and miR-146b-5p, both are well-established negative regulators of the NF-κB signaling pathway (2, 3, 20), supporting previous reports describing their role in attenuating inflammation and preventing progression to severe disease (6, 21, 22). In turn, MiR-223-5p, is highly expressed in myeloid cells and associated with a controlled inflammatory response, possibly mediated by neutrophils, and with the downregulation of IL-2Rα, a key molecule in the formation of regulatory T cells (16, 23, 24). In relation to MiR-30c, it has been implicated in the modulation of antiviral responses via the SOCS1/IFN-λ axis (25–27) and also participates in autophagy and is involved with oxidative stress, a hallmark of the miR-30 family, including miR-30c, in various biological contexts. Furthermore, its role extends to antigen presentation pathways (28–30). In turn, miR-144-3p is involved in erythropoiesis and oxidative stress regulation (31–33), whereas miR-126-3p plays a role in vascular integrity and angiogenesis, potentially offering endothelial protection in inflammatory settings (34). The upregulation of miR-126 observed in our cohort supports an early host endothelial and immune response in non-hospitalized patients, contrasting with the downregulation frequently described in late or severe stages of the disease, and reinforcing its potential as a biomarker for early detection of mild outcomes (35). Finally, miR-22-3p is a regulator of immune cell differentiation and inflammatory signaling (36).

Although miR-155 and miR-21-5p showed a tendency toward increased expression, these differences were not statistically significant. ROC curve analysis revealed limited discriminatory power for both miRNAs, with AUC values of 0.517 for miR-155 and 0.606 for miR-21-5p. These findings are different to earlier studies reporting upregulation of these miRNAs in COVID-19 patients. However, these findings were verified especially in severe cases (13, 20, 37–39), contrasting with our group (mild), demonstrating and reinforcing the differential expression of miRNA in different clinical presentations of the disease.

By applying ROC curve analysis to evaluate the ability of dysregulated miRNAs to act as biomarkers for COVID-19, heterogeneous diagnostic profiles were observed among the candidates. miR-30c demonstrated the highest individual diagnostic accuracy, with an AUC of 0.771, showing high sensitivity (100.0%) and moderate specificity (55.0%) at the Youden-optimal threshold, supporting its strong performance in detecting mild cases. Although direct evidence for miR-30c in COVID-19 is limited, its strong diagnostic performance has already been reported in other inflammatory context (40), reinforcing its potential analytical validity. miR-126 achieved an AUC of 0.763 with perfect specificity (100.0%) and moderate sensitivity (59.1%), suggesting suitability for confirmatory analysis where false positives must be minimized. Previous reviews have highlighted miR-126 as a potential COVID-19 biomarker due to its regulatory role in endothelial and inflammatory pathways (35, 37).

The miR-146b-5p, miR-223-5p, miR-144-3p, and miR-22-3p showed intermediate performance, with AUCs ranging from 0.684 to 0.719. miR-146b-5p reached an AUC of 0.719 (sensitivity 75.0%, specificity 66.7%), while miR-223-5p exhibited an AUC of 0.704 with high sensitivity (95.2%) but limited specificity (41.7%), indicating potential application in initial screening strategies (38). miR-144-3p (AUC = 0.685; sensitivity 66.7%, specificity 73.9%) and miR-22-3p (AUC = 0.684; sensitivity 94.4%, specificity 60.9%) also demonstrated moderate discriminatory performance. On the other hand, miR-146a presented an AUC of 0.702 with a sensitivity of 47.4% and a specificity of 87.5. Although miR-146a, miR-21-5p, and miR-155 presented varying individual diagnostic metrics in this study, all have been widely reported in the literature as dysregulated in COVID-19 patients. Garg et al. demonstrated increased serum levels of miR-21 and miR-155 in COVID-19 compared to healthy individuals (39). Furthermore, another study demonstrated that miR-155 achieved an AUC of 0.986, distinguishing COVID-19 patients from controls (41). However, the groups analyzed in these two studies were different, including severe or long COVID-19, respectively, suggesting that population context and patients’ characteristics may affect the analysis and must be considered.

It is important to highlight that although the curves exhibited different inclinations and step patterns, most deviated substantially from the reference line (AUC = 0.5), which represents random classification. These results indicate that the identified miRNAs are not only differentially expressed but also demonstrate potential utility for the diagnosis or screening of mild COVID-19 cases. Finally, these findings support the concept of “multiple miRNAs – one disease,” as opposed to the traditional “one miRNA – one disease” model (38), emphasizing that a panel of miRNAs with complementary sensitivity and specificity profiles may enhance overall diagnostic performance compared to individual biomarkers.

In relation to the predictive interaction network analysis (Figure 3), it indicated that the upregulated miRNAs target key genes within immune signaling pathways. hsa-miR-146a was predicted to target IRAK1 and TRAF6, both key mediators of Toll-like receptor and NF-κB signaling (42, 43), while hsa-miR-146b-5p was linked to STAT1, IRAK1, and TRAF6. hsa-miR-126-3p showed predicted interactions with CXCL12 (45), targets JAK1 (46), and DNMT1 (48, 49), as well as SOX2 within the broader target prediction dataset (47), genes implicated in cytokine signaling (50, 51), tissue repair (52, 53), and epigenetic regulation (49). hsa-miR-30c-5p was associated with NOTCH1 (54), BCL2L11 (55), CXCL12, and DNMT1, targeting pathways in adaptive immunity, cellular apoptosis, and methylation (54, 56–58). hsa-miR-144-3p shares several targets in this network, including SMAD4 (59, 60), NOTCH1 (61–63), and CXCL12, which are involved in TGF-β signaling (43, 44) and cell fate determination. In addition, hsa-miR-223-5p was linked to IL6R and BCL2L11, whereas hsa-miR-22-3p was predicted to interact with SIRT1, FOXO3, PTEN, and BCL2L11, which serve as crucial regulators of cellular stress response, metabolic homeostasis, and survival. These findings support the hypothesis that the observed miRNA signature may contribute to a coordinated host response aimed at controlling infection while preserving tissue homeostasis.

Additionally, several other predicted targets such as STAT1, IRAK1, TRAF6, SIRT1, FOXO3, and PTEN reinforce this hypothesis. STAT1 plays a critical role in the antiviral interferon response and is essential for the transcriptional activation of interferon-stimulated genes (64), while IRAK1 and TRAF6 are key signaling intermediates in Toll-like receptor and NF-κB pathways, both of which are central to early immune activation and cytokine production during viral infections (65–68). Their regulation by miR-146a and miR-146b-5p suggests a negative feedback loop controlling inflammatory responses (67). On the other hand, SIRT1 (69) and FOXO3 are regulators of cellular stress resistance, antioxidant responses, and autophagy, and may contribute to tissue protection under inflammatory conditions (70, 71). PTEN, a well-known tumor suppressor, also regulates immune homeostasis by modulating the PI3K–AKT pathway, thus influencing both innate and adaptive immunity (72). These additional targets further highlight the potential of the miRNA network to integrate immune activation with mechanisms that preserve epithelial integrity and prevent excessive inflammation.

Interestingly, functional enrichment analysis revealed a significant overlap between the predicted targets of the selected miRNAs and the transcriptomic signature denoted as RNA_Sun_Calu-3_12h_Up. This dataset refers to genes that were upregulated in Calu-3 human lung epithelial cells following 12 hours of SARS-CoV-2 infection, as reported by Sun et al. (19) These genes are involved in pathways consistently activated during viral infection, particularly type I interferon signaling, cytokine-mediated responses, and apoptosis regulation. The observed enrichment (–log₁₀P ≈ 5.2 for 12h and 3.6 for 24h) suggests that the miRNA-regulated gene set mimics or modulates host cellular responses during the early stages of infection. In this context, Calu-3 cells constitute a relevant in vitro model for studying SARS-CoV-2 pathogenesis due to their high permissiveness to viral entry and replication, as well as their ability to mount a robust innate immune response (19, 73, 74). The convergence of miRNA targets with this infection-induced expression profile supports the hypothesis that these miRNAs are functionally involved in the regulation of genes critical for viral sensing, cytokine signaling, and epithelial barrier function during SARS-CoV-2 infection (74). Taken together, these findings reinforce the biological plausibility of the miRNAs investigated in this study as modulators of key host responses in the respiratory tract, and potentially as biomarkers or therapeutic targets in COVID-19 (74, 75).

Altogether, these observations point toward a possible protective or modulatory role of the identified miRNAs in mild COVID-19; however, further functional validation is essential, as miRNA expressions are influenced by multiple factors, including viral load, host immune status, and genetic background (14, 76–78). In addition, the predicted interactions were derived from computational algorithms and require confirmation through in vitro and in vivo experiments. That way, longitudinal studies in larger and more diverse cohorts will be critical to elucidate the regulatory mechanisms mediated by these miRNAs and to validate their clinical utility as biomarkers (14). Finally, these upregulated miRNAs may simply be associated with the SARS-CoV-2 infection and not with outcome. To verify this, it is necessary to include comparison with severe cases. In any case, the upregulation of several miRNAs demonstrates a relevant difference between infected and uninfected individuals, and this information can be very useful for understanding viral infection and its modulation.

We identified an upregulated miRNA signature in patients with mild COVID-19, comprising miR-126, miR-146b-5p, miR-223-5p, miR-144-3p, miR-22, miR-146a, and miR-30c. The findings demonstrate the potential of these miRNAs as biomarkers for mild cases compared with uninfected individuals, with miR-30c showing the highest discriminatory power based on ROC curve analysis.

In addition, interaction network analysis identified predicted targets of the upregulated miRNAs, such as CXCL12, STAT1, IRAK1, TRAF6, SMAD4, JAK1, NOTCH1, and DNMT1, that are functionally associated with immune regulation, inflammation, and tissue remodeling. Complementarily, functional enrichment analysis performed with Metascape revealed the convergence of these miRNAs in key pathways related to innate immune responses, cytokine signaling, and tissue remodeling, further reinforcing the biological relevance of this signature. These findings suggest that the observed upregulation may not be a mere passive consequence of infection, but rather part of active host mechanisms aimed at modulating inflammation and immune responses. Such a miRNA-mediated ability to orchestrate a controlled and adaptive immune response may represent a key determinant in patients with favorable outcomes.

This study has several limitations that should be considered. First, the relatively small sample size (25 cases and 25 controls) limits the statistical power of our analysis, particularly for miRNAs that did not reach significance (e.g., miR-155 and miR-21-5p). A formal sample size calculation was not performed a priori, and these findings should be validated in larger, independent, and more diverse cohorts. Additionally, nine miRNAs were tested independently without correction for multiple comparisons. Consequently, some of the more marginal associations (e.g., miR-22, miR-144-3p) should be interpreted with appropriate caution until confirmed by orthogonal methods or replication. Second, this is a cross-sectional, single-timepoint study comparing individuals with mild/asymptomatic COVID-19 to uninfected controls. This design does not allow us to assess the diagnostic or prognostic value of these miRNAs across a spectrum of disease severity, nor their potential utility in disease monitoring, since no longitudinal sampling or comparison with moderate-to-severe cases was performed. The observed miRNA signature may therefore reflect general host response to SARS-CoV-2 infection rather than a severity- or outcome-specific biomarker profile, and future studies incorporating serial sampling and a broader severity spectrum are needed to clarify this distinction. Third, control status was defined by a single negative RT-qPCR result at the time of sample collection. Serological testing was not performed to rule out prior or subclinical SARS-CoV-2 infection, which could theoretically influence circulating miRNA levels independent of current infection status. Finally, given that RT-qPCR already provides high diagnostic accuracy for SARS-CoV-2, the clinical scenarios in which this miRNA panel would offer added value over existing molecular diagnostics (for example, as a complementary tool when RT-qPCR results are inconclusive, or as part of a broader host-response signature) warrant further investigation and prospective validation in independent, multicenter cohorts.

In conclusion, this study provides valuable insights into the potential role of miRNAs in SARS-CoV-2 infection associated with mild, non-hospitalized cases. The findings related to target genes could suggest the potential of miRNAs for future diagnostic and therapeutic strategies. However, given the modest sample size and the absence of a severe-disease comparator or independent validation cohort, these results should be regarded as hypothesis-generating, confirming their diagnostic or biological relevance will require validation in larger, independent, and clinically diverse populations.

## MATERIALS AND METHODS

### Study Design

This study was carried out during August 2021–2022 with individuals who underwent SARS-CoV-2 testing via RT-qPCR at the Laboratório de Farmacogenômica e Epidemiologia Molecular (LAFEM) of the Universidade Estadual de Santa Cruz (UESC). Nasopharyngeal swabs and peripheral venous blood samples were obtained from individuals who consented to participate in the study as volunteers. All participants provided written informed consent.

### Groups Definition

We included 25 individuals that tested positive for SARS-CoV-2 through RT-qPCR (Cases) and 25 that tested negative (Controls). Cases were non-hospitalized outpatients who presented at least one symptom at diagnosis (cough, sore throat, fever, rhinorrhea, anosmia, or ageusia), whereas controls were asymptomatic at the time of sampling.

### Laboratory Detection of SARS-CoV-2 Infection

Viral RNA extraction was performed using the automated Loccus EXTRACTA 32 system and the MVXA-P016 kit. SARS-CoV-2 RNA detection was carried out by RT-qPCR using the Allplex™ 2019-nCoV assay (Seegene®, Seoul, Korea) and the SARS-CoV-2 EDx assay (Bio-Manguinhos, FIOCRUZ, Brazil), following the manufacturers’ protocols. The 7500 Fast Real-Time PCR System (Applied BiosystemsTM, Life Technologies, USA), was used for amplification. Results were categorized as positive (SARS-CoV-2 detected), negative (SARS-CoV-2 not detected - uninfected) or inconclusive, with inconclusive cases excluded from further analysis.

### Sample processing, RNA Isolation, quantification, and differential expression

Blood samples were drawn using a 21-gauge needle (Becton-Dickinson, Franklin Lakes, NJ, USA) into tubes containing K2-EDTA. Plasma was separated by centrifugation, and 200 µL of the plasma sample was combined with 600 µL of Trizol LS (Invitrogen). The mixture was then stored at −80°C for up to 24 hours following blood collection. RNA isolation was performed by adding 200 µL of cold chloroform (∼8°C), followed by gentle inversion, incubation at room temperature for 5 minutes, and centrifugation at 14,000 rpm for 20 minutes at 4°C to separate the phases. The aqueous phase, averaging 600 µL in volume, was carefully transferred to a separate collection tube. Next, 1000 µL of absolute isopropanol was added, and the mixture was stored at −20°C for approximately 12 hours. After this period, the samples were thawed and centrifuged at 14,000 rpm for 20 minutes at 4°C. The supernatant was discarded, preserving the pellet for further analysis. Subsequently, 1,000 µL of 70% ethanol was added to the tube, followed by another centrifugation under the same conditions. The supernatant was again removed, and the pellet was retained. The tubes containing the pellets were left to dry at room temperature for approximately 15 minutes. Finally, the pellet was resuspended in 20 µL of RNase-free water and stored at −80°C. All reagents were maintained at approximately 8°C during the procedure.

RNA concentration was assessed using the NanoDrop 1000 spectrophotometer (Thermo Scientific). Only RNA samples with a 260/280 absorbance ratio of at least 1.8 were included in further analyses. Reverse transcription was performed using 500 ng of RNA with specific primers for microRNAs and the TaqMan MicroRNA Reverse Transcription Kit (Applied Biosystems) in a reaction volume of 20 µL, following the manufacturer’s instructions. The cDNA synthesis was conducted using the TaqMan MicroRNA Reverse Transcription Kit (Applied Biosystems) and stem-loop primers targeting, miR-155, miR-126-3p, miR-223-5p, miR-146a, miR-21-5p, miR-146b-5p, miR-144-3p, miR-22-3p and miR-30c-5p (Thermo Fisher assay IDs: 002287, 002228, 002098, 002163, 009397, 001097, 001097, 000398, 000419). All gene expression levels were normalized using the snU6 gene (Thermo Fisher assay ID: 001973). The reverse transcription reaction was carried out using 5 µL of purified microRNA in a total volume of 20 µL, following the recommended protocol: incubation at 16°C for 30 minutes, 42°C for 30 minutes, and 85°C for 5 minutes.

For qPCR analysis, each reaction contained 4.5 µL of cDNA, 0.5 µL of the TaqMan 20X Assay, and 10 µL of the universal PCR master mix (Applied Biosystems), in a final volume of 20 µL. Each sample was analyzed in duplicate, with thermal cycling conditions consisting of 95°C for 3 minutes, followed by 40 cycles of 95°C for 15 seconds and 60°C for 60 seconds, using a QuantStudio 3.0 system.

### Comparative analysis of miRNA expression

Normalization of miRNA expression was performed using the small nuclear RNA snU6, a commonly used endogenous control in miRNA studies, to minimize technical variability (79). Relative expression levels were calculated using the 2^−ΔΔCt method to compare target miRNA expression between groups. Outliers were identified and excluded using the ROUT method in GraphPad Prism version 11.0.2. Normality of the data was assessed using the Shapiro–Wilk test. Group comparisons were conducted using the Mann–Whitney U test. Associations between miRNA expression and clinical parameters were evaluated using Chi-square (χ²) and Fisher’s exact tests. All statistical analyses were performed in GraphPad Prism version 9, with significance set at p < 0.05.

### Receiver operating characteristic (ROC) curve analysis

The diagnostic performance of individual miRNAs was evaluated by receiver operating characteristic (ROC) curve analysis. ROC curves were generated by plotting the true positive rate (sensitivity) against the false positive rate (1 – specificity) at varying threshold settings. The area under the curve (AUC) was calculated to quantify the overall ability of each miRNA to discriminate between COVID-19 cases and controls. An AUC value of 0.5 indicates no discrimination, whereas an AUC of 1.0 reflects perfect discrimination. Sensitivity, specificity, and the optimal cutoff values were determined based on the Youden’s index (maximum [sensitivity + specificity – 1]). ROC analyses were performed using the GraphPad Prism (version 9.0). Confidence intervals (95% CI) for AUC values were computed to assess the precision of the estimates.

### Computational prediction of potential miRNA targets

Putative target genes of the differentially expressed miRNAs were predicted using the miRWalk 3.0 platform (http://mirwalk.umm.uni-heidelberg.de/) (80). This tool integrates multiple target prediction algorithms, including miRDB, TargetScan, and miRTarBase, providing a comprehensive dataset of miRNA-mRNA interactions. A stringent minimum score threshold of 0.95 was applied to enhance prediction accuracy. The resulting miRNA– mRNA interaction data were imported into Cytoscape version 3.10 for network construction and visualization (81). To explore the biological significance of the predicted targets, functional enrichment analysis was performed using the ClueGO plugin (version 2.5.10) (82), which integrates Gene Ontology (GO) terms and Reactome pathways (83). The bipartite miRNA–gene interaction network was constructed using the NetworkX Python package (version 3.11), connecting miRNAs to their experimentally validated target genes. Nodes were colored by type (miRNAs in orange, genes in light blue) with black labels for clarity. The layout was generated using the spring_layout algorithm (k = 0.8, seed = 42) to ensure reproducibility and visualize co-regulatory interactions among miRNAs sharing common targets.

### Statistical analysis

A Pearson’s chi-squared or Fisher’s exact test (with Bonferroni and Benjamini-Hochberg False Discovery Rate [FDR] corrections for multiple comparisons, respectively) with significance level of 5% (p ≤ 0.05) were applied to compare Case and Control groups. ROC curves were established for discriminating individuals with mild COVID-19 and uninfected individuals. The optimal sensitivity and specificity from ROC curves were determined by commonly used methods. All statistical calculations were performed by using GraphPad Prism software version 11.0.2 (GraphPad Software, La Jolla, CA).

## Data availability

The datasets generated during and/or analyzed during the current study are not publicly available but are available from the corresponding author on reasonable request.

## ETHICS APPROVAL

The study adhered to the guidelines outlined in the Declaration of Helsinki and received approval from the Research Ethics Committee of the State University of Santa Cruz under registration number CAAE: 38627420.3.0000.5526. Informed consent was obtained from all participants, and all data was handled confidentially and anonymized to ensure the privacy of the individuals involved.

## ACKNOWLEDGMENTS

We thank the Programa de Pós-graduação em Biologia e Biotecnologia de Microrganismos (PPGBBM) and the Programa de Pós-graduação em Ciências da Saúde (PPGCS).

